# Hyaluronic Acid on the Urokinase Sustained Release with a Hydrogel System Composed of Poloxamer 407

**DOI:** 10.1101/2019.12.31.891614

**Authors:** Hao-Ying Hsieh, Wei-Yang Lin, An Li Lee, Yi-Chen Li, Yi-Jane Chen, Ke-Cheng Chen, Tai-Horng Young

**Author notes:** Corresponding author information: Tai-Horng Young, PhD, Department of Biomedical Engineering, National Taiwan University, Taipei, Taiwan, #81455. These authors contributed equally to this work (Hao-Ying Hsieh, Wei-Yang Lin).

## Abstract

Pleural empyema is an inflammatory condition characterized by accumulation of pus inside the pleural cavity, which is usually followed by bacterial pneumonia. During the disease process, the pro-inflammatory and pro-fibrotic cytokines in the purulent pleural effusion cause proliferation of fibroblasts and deposition of extracellular matrix, which lead to fibrin deposition and fibrothorax. Urokinase instillation therapy through a chest drainage tube is frequently used for fibrinolysis in patients with empyema. However, urokinase treatment requires multiple instillation (2-3 times per day, for 4-8 days) and easily flows out from the chest drainage tube due to its high water solubility. In this in vitro study, we developed a thermo-responsive hydrogel based on poloxamer 407 (P407) combined with hyaluronic acid (HA) for optimal loading and release of urokinase. Our results show that the addition of HA to poloxamer gels provides a significantly more compact microstructure, with smaller pore sizes (**p < 0.001). The differential scanning calorimetry (DSC) profile revealed no influence on the micellization intensity of poloxamer gel by HA. The 25% poloxamer-based gel was significantly superior to the 23% poloxamer-based gel, with slower gel erosion when comparing the 16th hour residual gel weight of both gels (*p < 0.05; **p < 0.001). The 25% poloxamer-HA gel also exhibited a superior urokinase release profile and longer release time. A Fourier-transform infrared spectroscopy (FT-IR) study of the P407/HA hydrogel showed no chemical interactions between P407 and HA in the hydrogel system. The thermoresponsive P407/HA hydrogel may have a promising potential in the loading and delivery of hydrophilic drugs. On top of that, in vitro toxicity test of this combination demonstrates a lower toxicity.

## 1. Introduction

Pleural empyema is an inflammatory condition characterized by accumulation of pus inside the pleural cavity, which is usually followed by bacterial pneumonia [1]. Pleural effusion is found in 9-30% of cases of bacterial pneumonia, 20% of which progress to empyema. The pro-inflammatory and pro-fibrotic cytokines in the purulent pleural effusion cause the proliferation of fibroblasts and deposition of extracellular matrix, which lead to fibrin deposition and fibrothorax [2]. Urokinase instillation therapy through a chest drainage tube is a standard protocol used for fibrinolysis of the deposited fibrin material [3]. However, urokinase treatment requires multiple instillation (2-3 times per day, for 4-8 days), and easily flows out from the chest drainage tube due to its high water solubility [4, 5]. Therefore, we attempted to develop a thermo-responsive hydrogel based on poloxamer 407 (P407) combined with hyaluronic acid (HA) for the loading and release of urokinase and for over 24 hours.

P407 is a non-ionic tri-block copolymer composed of a central hydrophobic block of polypropylene oxide sandwiched between two hydrophilic blocks of polyethylene oxide (PEOx-PPOy-PEOx) [6]. P407 is a thermo-responsive polymer that is liquid at low temperatures; and the aqueous polymer solution converts to a gel status when the temperature rises. The solubility of the hydrophobic blocks decreases as they aggregate to minimize the interaction of PPO blocks with the solvent [7, 8]. Due to its thermo-responsive character, good biocompatibility, and low toxicity [9], P407 is widely used for smart drug delivery [8] and in different formulations such as nasal, ophthalmic, and vaginal [10-14]. When used on its own, the P407 gel rapidly loses its gelation ability after dilution in a water-rich environment. Blending P407 with other polymers or molecules for better drug loading and retention has been previously studied [7, 8, 15]. For example, the P407 based hydrogels have been widely used to encapsulate some small molecular drugs (with a molecular weight (MW) under 500g/mole) such as ketorolac, metoprolol, and doxycycline; and the P407-based hydrogels could continually release these drugs for up to 20 hours [15, 16]. However, urokinase is a hydrophilic macromolecular protein kinase with 411 amino acid residues and a molecular weight of 32 kDa [17, 18], which facilitates the movement of water molecules into the urokinase-loaded P407 gel and therefore faster dissolution of P407-based hydrogels due to the larger constitutional property of urokinase, as shown in a previously reported *in vivo* study [19]. Therefore, a proper additive is therefore needed in order to achieve an optimized release of urokinase with the P407-based gel.

Hyaluronic acid (HA) is an endogenous glycosaminoglycan that exists in various tissues, including connective tissues and the aqueous humor of the eye. This polysaccharide is composed of disaccharide monomers (N-acetylglucosamine and glucuronic acid) and plays an important role in tissue hydration, cell migration, and wound healing [20]. HA is highly hydrophilic and able to form hydrogen bonds with water molecules. Also, HA was used as a drug delivery agent with different routes of administration in ophthalmic, nasal, pulmonary and oral [21], which has been reported to have the improved rheological and mucoadhesive properties [22, 23]. In addition to the pure HA, HA/Poloxamer hydrogels were studied as a thermo-responsive injectable filling material with the capability of controlled drug release for bone regeneration [24]. However, those studies were devoted to chemical synthesis and formulations. The release kinetics of urokinase and the effect of HA on urokinase-loaded drug release system have not been reported yet. In this study, we investigated different combinations of P407 and HA for the optimal effect of urokinase loading and release. Micellization and gelation behavior, gel dissolution, release of urokinase and its stability within the gels, and gel microstructure were also studied.

## 2. Materials and Methods

### 2.1. Materials

In this study, we aimed to reduce the frequency to single administration daily at maximal in vivo regimen release. Poloxamer 407 (P407, culture tested) was obtained from Sigma-Aldrich (Gillingham, UK). Research-grade sodium hyaluronate (HA) of 490±11.3 kDa was purchased from Lifecore Biomedical, Inc. (Minnesota, USA). Lyophilized urokinase powder for injection (60000 IU) was purchased from Taiwan Green Cross Co., Ltd (Taiwan).

### 2.2. Sample preparation

23% and 25% w/w P407 aqueous solution was prepared using a ‘cold method’ [7, 8]. Briefly, a weighed amount of P407 was added to water that had been equilibrated at 4–8 °C before use. The poloxamer solution was then kept for a further 24–36 hours in a refrigerator until a clear solution was obtained. Under an ice bath (temperature maintained at 2-5 °C), 5, 6, or 7 mg of HA was added into every ml of 23% or 25% cold P407 aqueous solution for dissolution under gentle stirring for 2 h. Then, 60000 IU lyophilized urokinase was dissolved into 60 mL water to make a 1000 IU/mL aqueous solution. 100 IU urokinase (0.1 ml of 1000 IU/ml urokinase aqueous solution) was loaded into 1 mL of P407/HA solution on ice bath with gently stirred.

### 2.3. Fourier transform infrared spectroscopy (FT-IR)

Fourier-transform infrared spectroscopy (FT-IR) experiments were performed using a Spectrum 100 spectrometer (PerkinElmer, USA). The samples were previously ground and mixed thoroughly with potassium bromide (1 mg of sample to 80 mg of potassium bromide). Potassium bromide discs were prepared by compressing the powders in a hydraulic press. Scans were obtained at a resolution of 1 cm^−1^ from 4000 to 450 cm^−1^.

### 2.4. Gel dissolution

Dissolution profiles of the P407-based gels in an aqueous environment were determined using the gravimetric method [25]. A pre-weighed glass vial of 13 mm diameter containing 0.6 g of the gel was equilibrated at 37 °C, and 0.3 mL of water previously equilibrated at 37°C was layered over it. The liquid medium was removed at pre-determined time intervals, the vial was re-weighed, and the weight of residual gel was calculated from the difference in weight of the vial. The entire process was carried out in an incubation room maintained at 37 °C.

In vitro cumulative release of urokinase

A membrane-less dissolution model was applied to the study of the release profile of urokinase [25]. The urokinase-loaded gels were treated as described above. The concentration of urokinase in the released medium was collected at pre-determined time intervals, and then determined using a fluorometric spectrophotometer utilizing the Abcam^®^ Urokinase Fluorometric Activity Assay Kit (Abcam, USA). An AMC-based peptide substrate containing the recognition sequence for urokinase was used. The urokinase present in the sample catalyzes the cleavage of the substrate and releases AMC, which could be easily quantified by measuring its fluorescence at Ex/Em = 350/450 nm with an ELISA (SpectraMax M2, Molecular Devices, USA).

### 2.5. Differential scanning calorimetry (DSC)

Calorimetry was carried out using a MicroCal VP-DSC microcalorimeter (Malvern, USA) equipped with VP Viewer software. Approximately 5 mg of poloxamer aqueous solution was placed in sealed aluminum pans. Prior to measurement, the sample was subjected to the following thermal cycle: heating from 0 to 50 °C, then cooling from 50 to 0 °C at a rate of 10 °C/min. DSC traces were then recorded as the temperature increased from 0 to 50°C at a rate of 10 °C/min, with an empty pan as a reference. Data were analyzed to obtain onset temperature (T_onset_), area under the peak, peak temperature (T_peak_), and endset temperature (T_endset_) of the endothermic peak.

### 2.6. Scanning electron microscope (SEM)

The samples were processed through lyophilization, which was used to preserve the cross-sections of samples. The samples were then coated with gold before being placed onto SEM specimen holders. The microstructures were observed under an S-4800 scanning electron microscope (Hitachi, Japan).

### 2.7. In vitro cytotoxicity

The cytotoxicity of 23% and 25% P407, urokinase and sodium hyaluronate were measured by Alamar blue assay. Normal fibroblasts (Hs68 cells) were seeded at a density of 2×10^4 (cell/ml) per well in 1ml culture medium (DMEM-HG) supplemented with 10% FBS in a 48-well plate and incubated for 3 hours at 37°C in a humidified incubator containing 5% CO_2_. Urokinase concentrations tested were 1000(IU/ml)[26]. The P407: urokinase proportion was maintained at 9:1(ml) for all formulations into the cell culture wells, at final volume of 1 ml. Subsequently the Alamar blue fluorescence was quantified at excitation and emission wavelength of 560nm and 590nm respectively by ELISA plate reader. For each plate the reading was done in quadruplicate (n=4)[27].

## Results

### 3.1. FT-IR spectroscopy characterization

The FT-IR spectra of P407, HA, urokinase, and their physical mixtures (P407 + urokinase and P407 + HA + urokinase) were performed in order to investigate the molecular interactions between P407, HA, and urokinase. The P407 spectra revealed characteristic peaks of CH_2_ stretch (2920 cm^−1^), C-O stretch (1100 cm^−1^), and C-O-C linkage (952 cm^−1^). This was comparable with the previously recorded FT-IR spectra of P407 [16]. The HA spectra showed an absorption peak at 1732 cm^−1^, corresponding to a C=O stretch. The spectra of the physical mixture (P407 + HA + urokinase) showed all the characteristic peaks of each chemicals without any shifts (Figure 1). Based on the FT-IR spectra, no chemical interactions between drug and polymer were observed in the physical mixtures.

**Figure 1.**
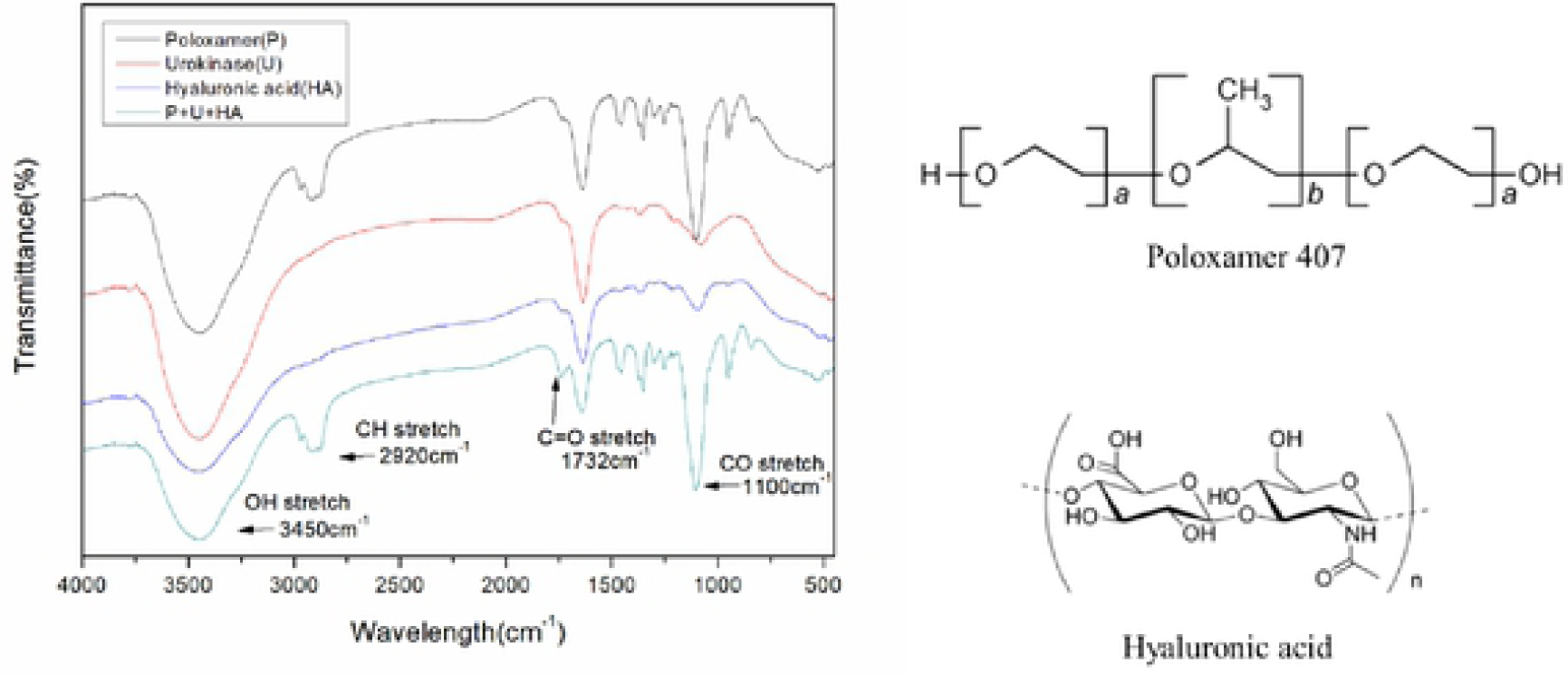
FT-IR spectra **(a)** of aqueous P407, U, P407+U, HA, and P407+U+HA. Chemical structure of **(b)** P407 and **(c)** HA.

### 3.2. Differential scanning calorimetry (DSC)

A DSC profile was used to determine the micellization behavior of P407 gels. The illustrative thermogram showed endothermic peaks indicative of micellization. The micellization process is characterized by the onset temperature (T_onset_), the area under the peak that reflects the enthalpy change, the peak temperature (T_peak_) [28], and the endset temperature (T_endset_) [29]. T_onset_ is the temperature at which micelles begin to form, while T_endset_ is the temperature at which the micellization process is completed. The Tpeak is referred to as the critical micellization temperature (CMT). The endothermic peak arises from the dehydration of the relatively hydrophobic polypropylene oxide (PPO) blocks of the P407 molecules during the micellization process, as previously reported [30].

The DSC thermogram showed a decrease in CMT when HA was added into pure 23% and 25% aqueous P407 solution (Figure 2). The micellization process occurred earlier in all aqueous P407/HA solutions as shown by the decrease in T_onset_. The effect of added HA on micellization intensity is relatively minor. On the other hand, the addition of urokinase into aqueous P407 solution delayed the micellization process with a markedly decrease in the micellization intensity. This is demonstrated by the increase in T_onset_, CMT, and T_endset_, as well as the decrease in the endothermic peak (Figure 2c).

**Figure 2.**
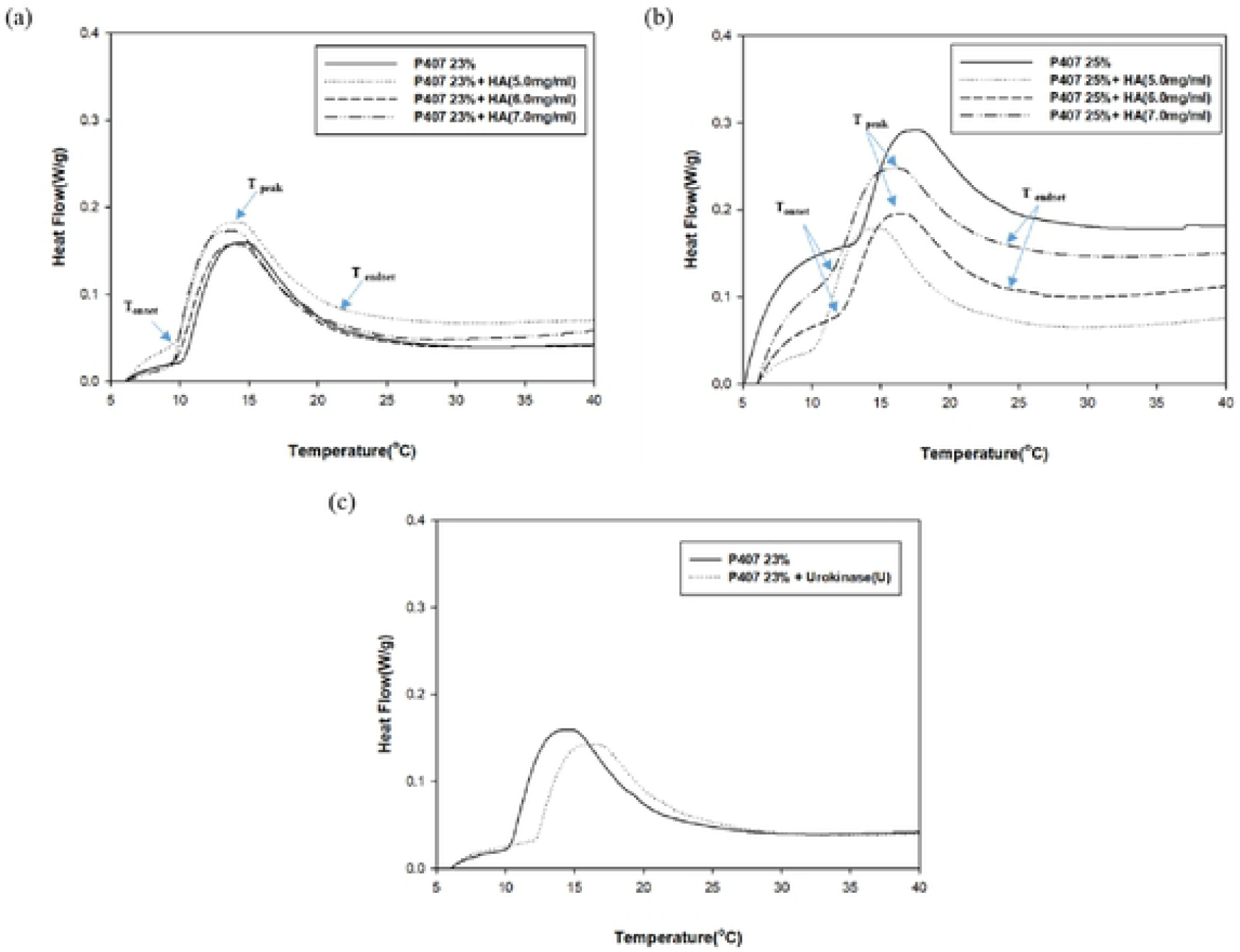
Micellization intensity P407-based gels. Tonset, CMT, and Tendset decreased when HA was added into the aqueous **(a)** 23% and **(b)** 25% P407. The influence of HA on micellization intensity was relatively minor. The micellization process was delayed and micellization intensity was decreased when **(c)** urokinase was added into aqueous P407 solution.

### 3.3. Dissolution of poloxamer gels

We examined the gel dissolution properties of pure P407 gels, P407 with HA, and P407 with HA and urokinase. The dissolution curve revealed no swelling of pure 23% P407 gel and a minor degree of swelling (3.2 ± 1.7%) of pure 25% P407 gel in the first hour. When HA was added into the P407 gels, a relatively wide degree of swelling ranging from 5.1 ± 1.7% to 17.5 ± 4.7% were observed in the first 3 hours. The degree of swelling increased with the increase of concentration of P407(23%, 25%) and HA (5.0mg/ml, 6.0mg/ml and 7.0mg/ml accordingly)(Figure 3a and b). The 25% P407 with 5mg/ml HA exhibited the highest degree of swelling (17.5 ± 4.7%).The addition of HA remarkably improved the hydrogel properties of pure P407 gels with extended gel dissolution time. When comparing the residual gel weights at 20^th^ hour)(Figure 3c), the addition of HA significantly slowed down the gel erosion process in the 23% P407 gels. However, no statistically significant difference was observed between the 25% P407 and 25% P407+HA when comparing the residual gel weight at 24^th^ hour)(Figure 3d).

**Figure 3.**
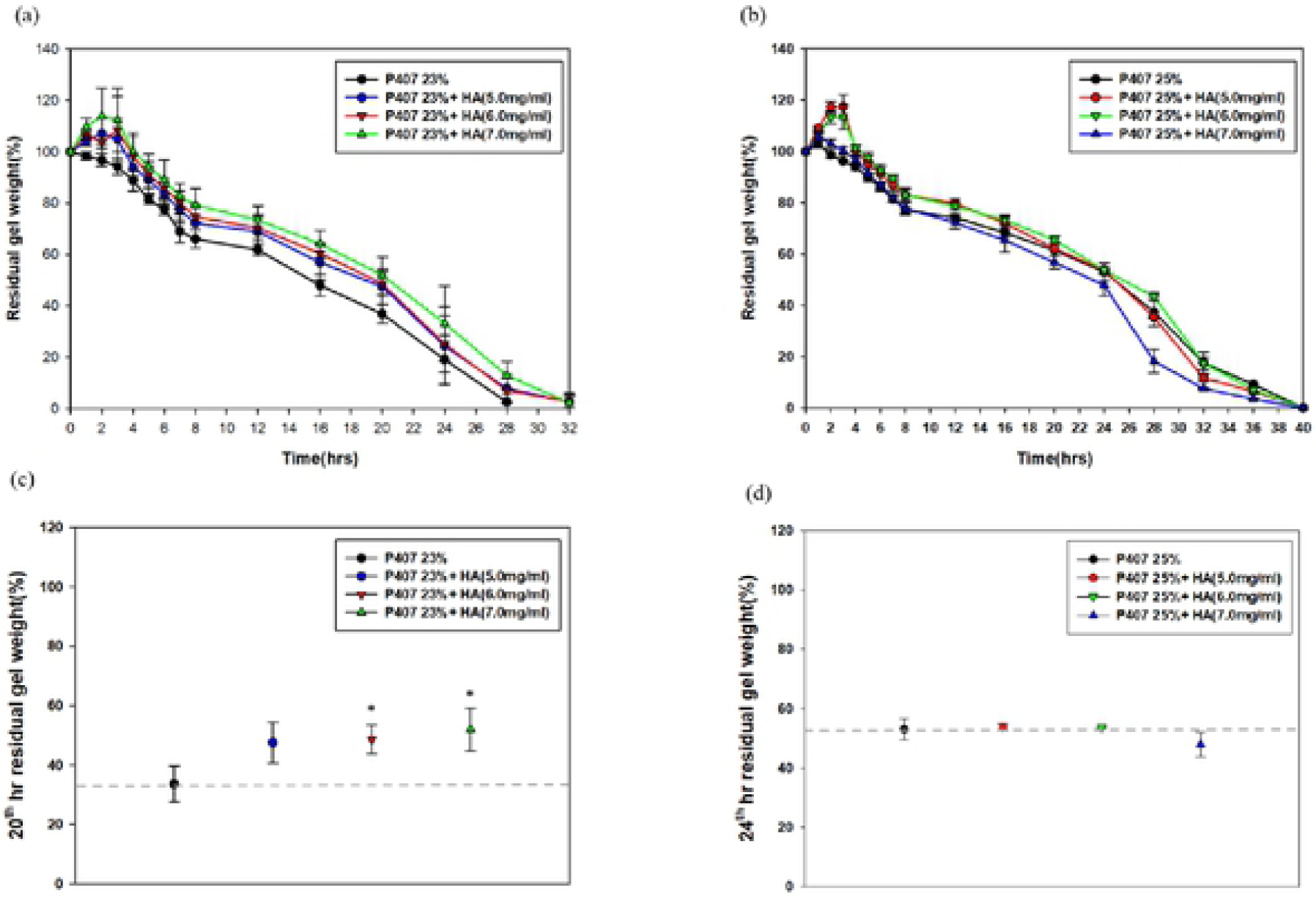
Gel dissolution profiles of **(a)** 23% and **(b)** 25% P407 with and without HA. They axis represents residual gel weight percentage at different time points. The addition of HA into 23% and 25% P407 gels markedly increases swelling rate in the first 3 hours. The addition of HA significantly slowed down the gel erosion in 23% and 25% P407 gels when compared at **(c)** 20^th^ and **(d)** 24^th^ hour, respectively. Significant differences from control group (P407+U) were denoted with \**P* < 0.05 and \*\**P* < 0.01.

Furthermore, the loading of urokinase into P407 gels accelerated the gel dissolution by 8-12 hours compared to pure P407 (Figures 4a and b). The addition of HA remarkably extended the gel erosion process in the urokinase-loaded 23% P407 when comparing the residual gel weights at 16^th^ hour. Similarly, the extended gel erosion process was observed in the 25% P407 +HA+urokinase at 20^th^ hour.

**Figure 4.**
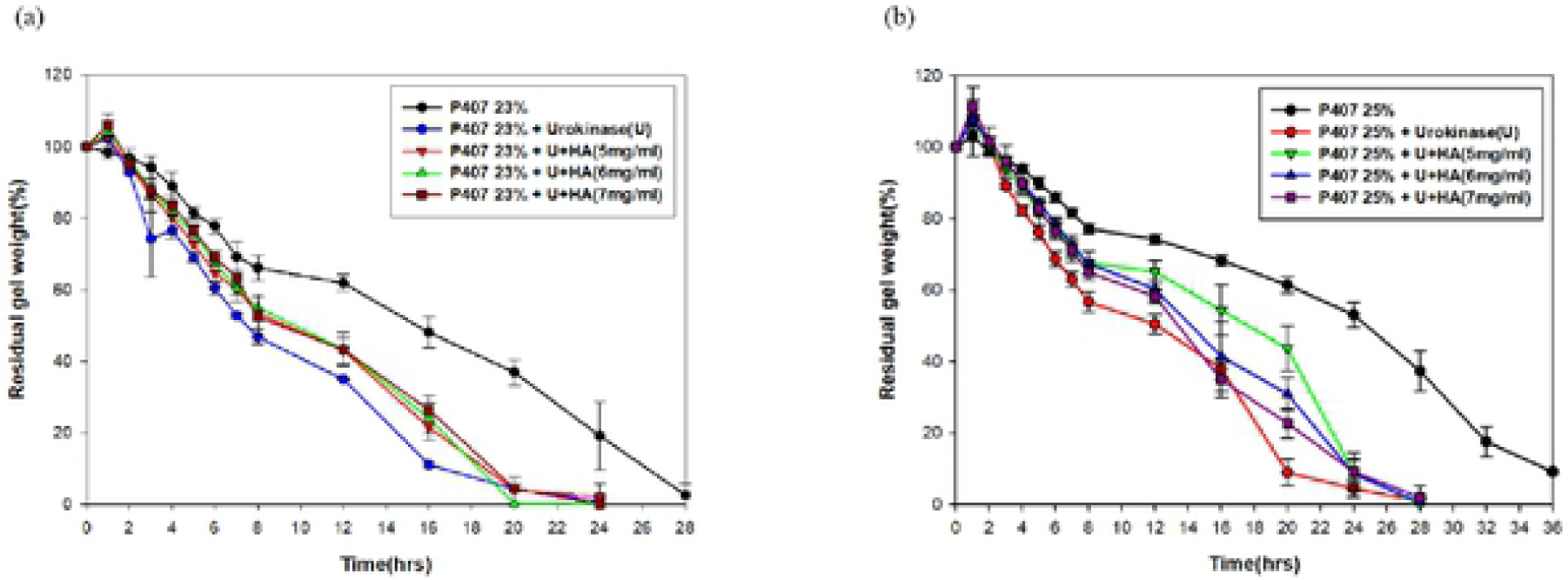
Gel dissolution profiles of **(a)** 23% and **(b)** 25% urokinase-loaded P407 with and without HA. The gel dissolution time decreased markedly in urokinase-loaded 23% and 25% P407 in the comparison with the non-urokinase-loaded gels.

### 3.4. Release of urokinase from P407 gels

The release profiles of urokinase in the 23% P407 and 25% P407 was found to last for 20 hours and 24 hours, respectively (Figure 5). Moreover, the release of urokinase within the gels had been augmented in a significant rate by the addition of HA, which was reinforced in the scenario of 25% P407.

**Figure 5.**
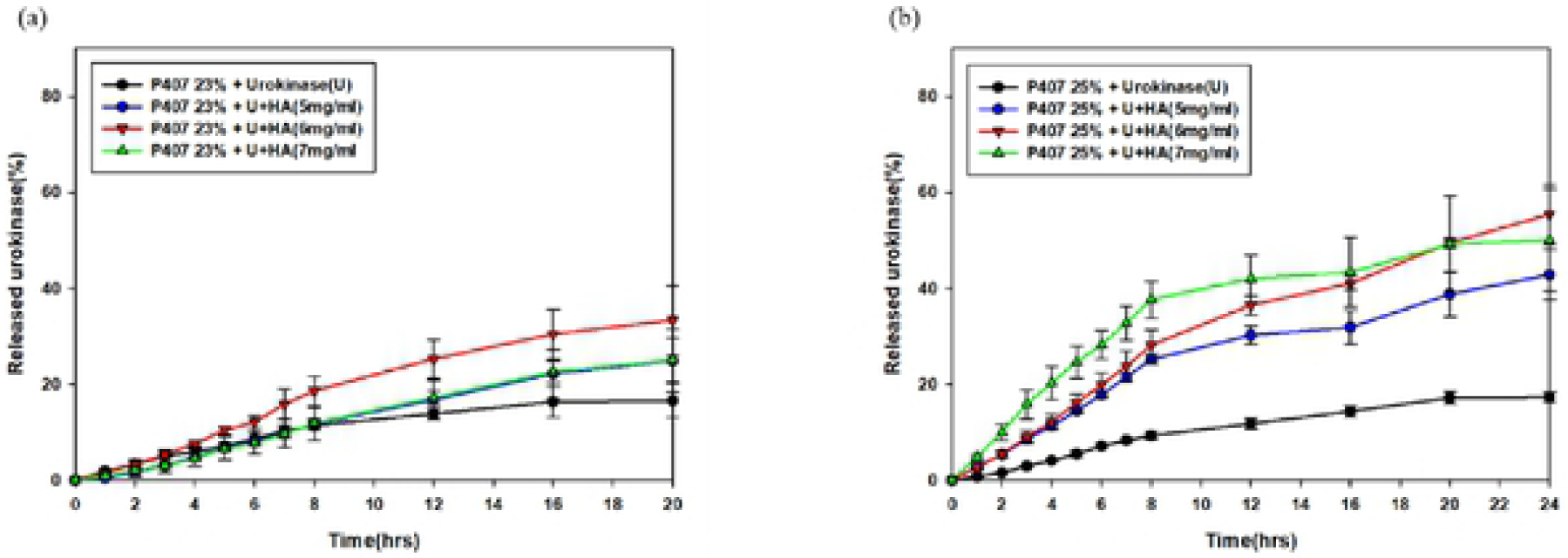
Urokinase release curves of 23% **(a)** and 25% **(b)** P407-based gels. The addition of HA into 25% P407 gel significantly increased urokinase activity.

### 3.5. Microstructure of P407 gels influenced by the addition of urokinase and HA

SEM images showed the structures of pure P407 gels as porous and sponge-like. When urokinase was loaded into the P407 gels, a significant change in the structure and pore size was observed. However, the addition of HA significantly reduced the pore size and resulted in a more compact porous structure (Figures 6), suggesting the possible protective role of HA in this hydrogel system. The changed microstructural may explain the extended gel erosion we have observed, hence resulting in the higher stability of urokinase at 37°C.

### 3.6. In vitro cytotoxicity

The correlation between cytotoxicity and individual urokinase, sodium hyaluronate, (Figures 7a)23% P407, (Figures 7b)25% P407 and their combination were assessed. The results showed that the cellular activities of above materials were above 80% of the control.

**Figure 7.**
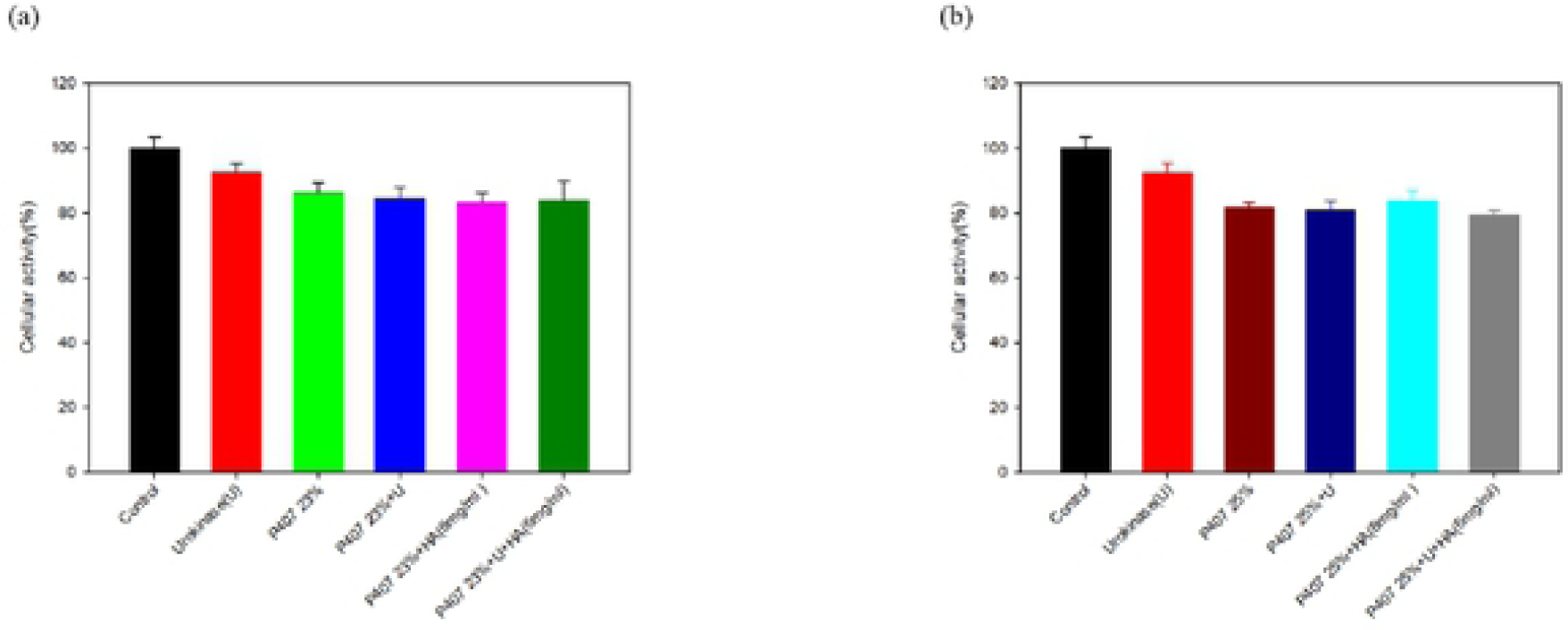
Alamar blue assay was used to determine the Hs68 cells cell viability. Urokinase and 6mg HA were added to (a) 23% P407 (b) 25% 407.

## 4. Discussion

P407, one of the members of the poly(PEOx-PPOy-PEOx) tri-block copolymers family, presents with a polyoxypropylene (PPO) molecular mass of 4,000 g/mol and a 70% polyoxyethylene (PEO) content. Both of the PEO and PPO blocks develop hydrogen bonds with water molecules, but the former is more hydrophilic than the latter. P407 is characterized by its temperature dependent self-assembling and thermogelling behavior. As the temperature rises, the aqueous P407 solution converts to a gel status driven by the decline of the solubility of PPO blocks, in which aggregation occurs in order to minimize their interaction with water molecules [7, 8]. The aggregated hydrophobic blocks comprise the core of the micelle, whilst the PEO blocks form the outer layer of hydrophilic shell that is bound to the water molecules covalently. The solution-to-gel transformation temperature is concentration-dependent and decreases as the concentration of P407 aqueous solution increases[30]. When the concentration is lower than 15% w/w, gel formation is absent regardless of the temperature change[19].

Large amount of micelles and aqueous channels make up poloxamer-based gels, making them excellent candidates to load hydrophilic molecules in spaces between micelles and channels. When the concentration of P407 is increased, a shorter inter-micellar distance and a greater tortuosity can be generated in the aqueous phase of the gel structure, resulting in more cross-links between micelles and a greater number of micelles per unit volume[31]. In spite of that, previous *in vivo* studies have explored the applicability of poloxamer-based gels and sustained delivery of model drug with the molecular weight of 5, 20, and 40 kDa. Their results implied that there is an association between high molecular weight of loaded hydrophilic drugs and faster gel erosion [19].

In our study, we observed an acceleration of gel erosion when urokinase was loaded into P407 gel. However, the addition of HA not only extended the gel dissolution time but also protected the activity of urokinase within gels, bringing forth a higher percentage of urokinase release. The acceleration of gel erosion might be attributed to the decrease in micellization intensity in the urokinase-loaded P407 gels. Adding HA into pure P407 gels also gave rise to a reduction in CMT without altering the intensity of micellization. The decrease in CMT and accelerated micellization process may represent the decrease in polarity and proportions of the hydrated methyl group in the PPO blocks[30, 32, 33]. In the context of hydrophilic drugs, gel erosion and drug diffusion occur simultaneously, which means that slower gel erosion often represents slower drug release [19, 31]. The results from our study concur with this phenomenon. In addition, our finding indicates that the existence of urokinase may result in the increase of the solubility of PPO blocks in aqueous P407 solution, possibly through formation of hydrogen bonds with the PPO blocks[34].

The addition of HA significantly postponed the event of gel erosion in the 23% P407 gels with a swelling remarkably observed. The increase in swelling of the hydrogel may occur on account of the COO^−^ groups within the HA/P407 hydrogels which induced the repulsive force, causing the infiltration of water infiltration and an extended process of gel erosion[24]. Although *in vitro* gel dissolution studies showed no significant increase in gel dissolution time regarding the addition of HA into the 25% P407, the swelling may not be the sole determining factor of the process of gel erosion[35-37]. We reasoned that this phenomenon is less prominent as that in the 25% P407 gels owing to the strong inter-micellar interaction associated with the increase of P407 concentration[36].

The microstructure of poloxamer gels revealed by SEM showed that urokinase induced a remarkable structural change with larger pore size. However, a more compact structure with smaller pore size was observed on the HA-added poloxamer gel, which may reflect the strong interaction between HA and poloxamer micelles. The compact porous structure also explains the high swelling rate of P407/HA gels which was observed in the first 3 hours during the gel dissolution[38]. This finding not only provides a good explanation of the slower erosion of P407/HA gels, but also reveals the strong physical interactions between the two polymers[39]. Negligible cell toxicity of in vitro toxicity test was detected when 23%, 25% P407 and urokinase were used.[40].

## 4. Conclusions

Our study presents a hydrogel system composed of P407 and HA that successfully achieved sustained release of urokinase for 24 hours. Sodium hyaluronate, as an additive into P407 gels, can decrease the CMT and accelerate the micellization process without affecting micellization intensity and CGT. The addition of HA was found to perform better on slowing down gel erosion and improving hydrogel property when mixed with aqueous P407 solution, hence resulting in a better regimen for longer urokinase release. HA also provides P407 hydrogel a higher swelling property and a more compact microstructure with smaller pore size. The P407/HA hydrogel is therefore a promising material for loading hydrophilic drugs in future given its safe property without causing cell toxicity after local injection.

## Acknowledgements

The authors declare that they have no conflicts of interest. The authors want to express their gratitude to the Department of Thoracic Surgery, Lo-Hsu Medical Foundation, Lotung Poh-Ai Hospital, Lotung, Taiwan and Ministry of Science and Technology, Taiwan (No.108-2314-B-002-117-MY3) for providing funding. The authors also acknowledge the technical assistance of the staff at the Eighth Core Lab of National Taiwan University Hospital. And the instrument supported from the department of electron microscopy, School of Science, Taiwan University, Ya-Yun Yang and Ching-Yen Lin.

